# Dissecting super-enhancer hierarchy based on chromatin interactions

**DOI:** 10.1101/149583

**Authors:** Jialiang Huang, Kailong Li, Wenqing Cai, Xin Liu, Yuannyu Zhang, Stuart H. Orkin, Jian Xu, Guo-Cheng Yuan

## Abstract

Recent studies have highlighted super-enhancers (SEs) as important regulatory elements for gene expression, but their intrinsic properties remain incompletely characterized. Through an integrative analysis of Hi-C and ChIP-seq data, we find that a significant fraction of SEs are hierarchically organized, containing both hub and non-hub enhancers. Hub enhancers share similar histone marks with non-hub enhancers, but are distinctly associated with cohesin and CTCF binding sites and disease-associated genetic variants. Genetic ablation of hub enhancers results in profound defects in gene activation and local chromatin landscape. As such, hub enhancers are the major constituents responsible for SE functional and structural organization.

## Introduction

Enhancers are *cis*-acting DNA sequences that control cell-type specific gene expression (Banerji, Rusconi, & Schaffner, 1981). Super-enhancers (SEs) are putative enhancer clusters with unusually high levels of enhancer activity and enrichment of enhancer-associated chromatin features including occupancy of master regulators, coactivators, Mediators and chromatin factors (Hnisz et al., 2013; Parker et al., 2013; Whyte et al., 2013). SEs are often in close proximity to critical cell identity-associated genes, supporting a model in which a small set of lineage-defining SEs determine cell identity in development and disease.

Despite the proposed prominent roles, the structural and functional differences between SEs and regular enhancers (REs) remain poorly understood (Pott & Lieb, 2015). A few SEs have been dissected by genetic manipulation of individual constituent enhancers. In some studies, the results are consistent with a model whereby SEs are composed of a hierarchy of both essential and dispensable constituent enhancers to coordinate gene transcription (Hay et al., 2016; Hnisz et al., 2015; Huang et al., 2016; H. Y. Shin et al., 2016). Due to the technical challenges in systematic characterization of SEs on a larger scale, it remains difficult to evaluate the generality of hierarchical SE organization in the mammalian genome.

Enhancer activities are mediated by the 3D chromatin interactions. Recent advances in Hi-C (Lieberman-Aiden et al., 2009) and ChIA-PET (Fullwood et al., 2009) technologies enable systematic interrogation of the genome-wide landscapes of chromatin interactions across multiple cell types and growth conditions (Dixon et al., 2015; Dixon et al., 2012; Dowen et al., 2014; Javierre et al., 2016; Ji et al., 2016; Jin et al., 2013; Rao et al., 2014; Tang et al., 2015). These data strongly indicate that the 3D chromatin organization is highly modular, containing compartments, topologically associating domains (TADs), and insulated neighborhoods. Of note, genomic loci with high frequency of chromatin interactions are highly enriched for SEs (Huang, Marco, Pinello, & Yuan, 2015; Schmitt et al., 2016), suggesting that proper 3D chromatin configuration may be essential for orchestrating SE activities.

Here we developed an approach to dissect the compositional organization of SEs, based on the patterns of long-range chromatin interactions. We found that a subset of SEs exhibits a hierarchical structure, and hub enhancers within hierarchical SEs play distinct roles in chromatin organization and gene activation. Our findings also identified a critical role for CTCF in organizing the structural (and hence functional) hierarchy of SEs.

## Results

### A subset of SEs contains hierarchical structure

To systematically characterize the structural organization of SEs, we developed a computational approach that integrates high resolution Hi-C and ChIP-seq data (Figure 1A), We defined SEs with the standard ROSE algorithm (Loven et al., 2013; Whyte et al., 2013). Briefly, neighboring enhancer elements defined based on H3K27ac ChIP-seq peaks were merged and ranked based on the H3K27ac ChIP-seq signal, where top ranked regions were designated as SEs. To quantify the degree of structural hierarchy associated with each SE, we defined a computational metric, called hierarchical score (or H-score for short), as follows. First, we divided each SE into 5kb bins to match the resolution of Hi-C data (Figure 1B). Next, we normalized the frequency of chromatin interactions within each SE by transforming the raw frequency values to z-scores. Third, we evaluated the maximum z-score across all bins in each SE, and referred to the outcome as the H-score associated with the SE. A higher H-score value indicates the chromatin interactions associated with a SE are mediated through a small subset of constitutive elements (Figure 1B). Fourth, by applying a threshold value of H-score, we divided all SEs into two categories, which we referred as hierarchical and non-hierarchical SEs, respectively (Figure 1B). Finally, if an enhancer element within hierarchical SEs is associated with a z-score greater than the threshold of H-score, the element is referred as a hub enhancer, whereas the remaining enhancers within the same SE are termed non-hub enhancers (Figure 1B).

**Figure 1.**
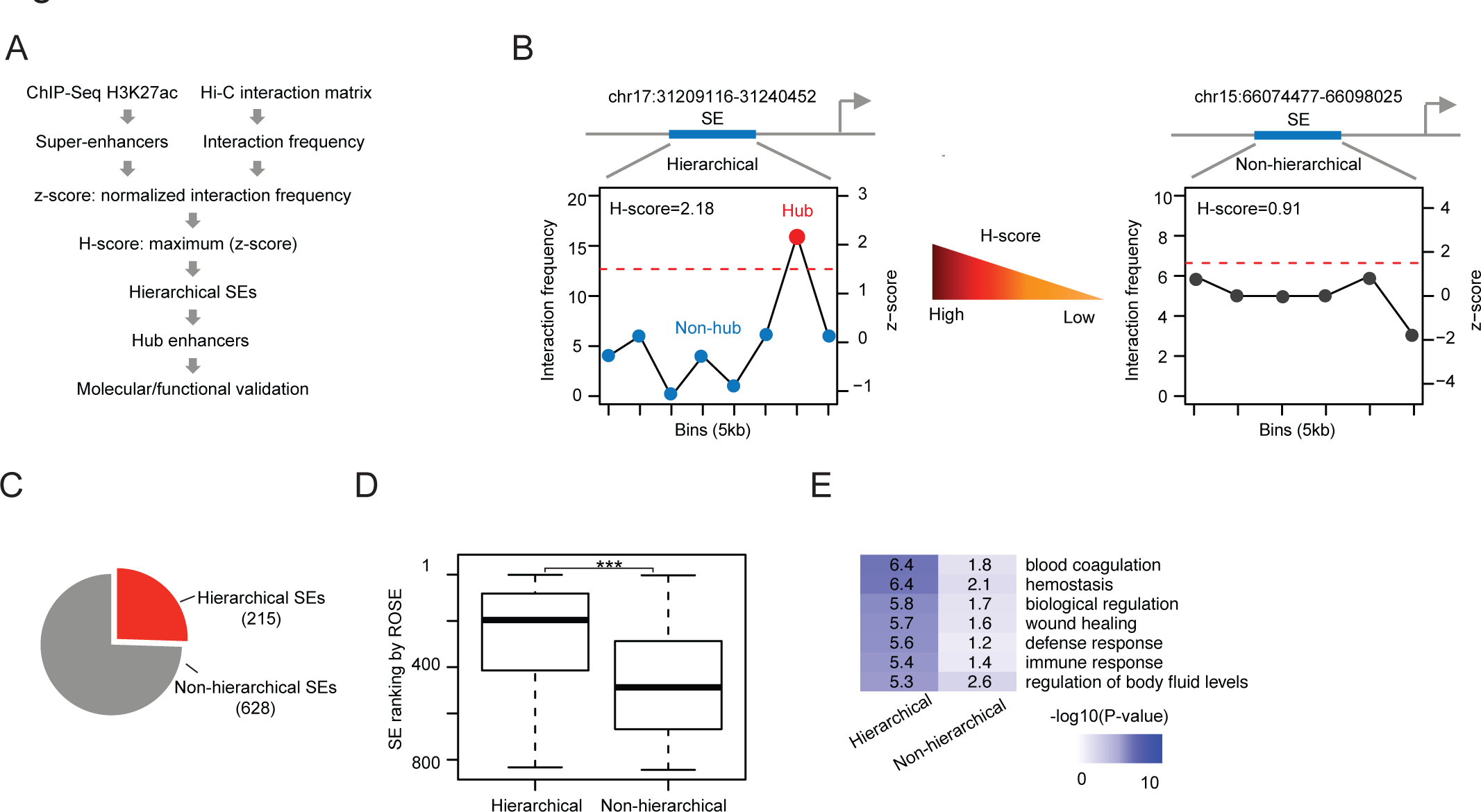
Definition of hierarchical SEs and hub enhancers based on Hi-C chromatin interactions. (**A**) Overview of pipeline. (**B**) Representative hierarchical (left) and non-hierarchical (right) SEs. For each 5kb bin within SE, the frequency of chromatin interactions (left *y-axis*) of and the z-score (right *y-axis*) are shown. The dashed red line represents the threshold of z-score = 1.5. (**C**) The proportion of hierarchical and non-hierarchical SEs. (**D**) The ROSE ranking of hierarchical and non-hierarchical SEs. *P* value is calculated using Wilcoxon rank-sum test. \**P* < 0.05; \*\**P* < 0.01; \*\*\**P* < 0.001. (**E**) GREAT functional analysis of hierarchical and non-hierarchical SEs.

We applied this pipeline to dissect the SE hierarchy in two human cell lines K562 (erythroleukemia cells) and GM12878 (B-cell lymphoblastoid cells), using publicly available high-resolution Hi-C and ChIP-seq data (T. E. P. Consortium, 2012; Rao et al., 2014). In total, we identified 843 and 834 SEs in K562 and GM12878 cells, respectively. On comparison of high-resolution (5kb) Hi-C profiles in K562 and GM12878 cells(Jin et al., 2013), we observed that SEs contain a significantly higher frequency of chromatin interactions than regular enhancers (*P* = 1.2E-69 in K562, *P* = 2.0E-123 in GM12878, Student’s t-test, Figure 1-figure supplement 1A), consistent with previous studies (Huang et al., 2015; Schmitt et al., 2016). By applying a threshold value of H-score = 1.5, which roughly corresponds to the 95^th^ percentile of z-scores (Figure 1-figure supplement 1B), we divided SEs into two categories: hierarchical and non-hierarchical SEs (Figure 1-figure supplement 1C). As expected, hub enhancers display a higher frequency of chromatin interactions than non-hub enhancers (Figure 1-figure supplement 1D). On average, both hub and non-hub enhancers within SEs contain a higher frequency of chromatin interactions than REs.

In total, we identified 215 (23% of all SEs) and 319 hierarchical SEs (34%) in K562 and GM12878 cells, respectively (Figure 1C and Figure 1-figure supplement 2A). The hierarchical SEs tend to be ranked higher than non-hierarchical SEs based by the ROSE algorithm (*P* = 1.2E-25 in K562, *P* = 2.5E-21 in GM12878, Wilcoxon rank-sum test, respectively, Figure 1D and Figure 1-figure supplement 2B). Using GREAT functional analysis (McLean et al., 2010), we observed that, compared with non-hierarchical SEs, hierarchical SEs were more enriched with gene ontology (GO) terms associated with cell-type-specific biological processes, such as ‘blood coagulation’ in K562 cells and ‘B cell homeostasis’ in GM12878 cells (Figure 1E and Figure 1-figure supplement 2C). These results suggest that hierarchical SEs may play a more important role in the maintenance of cell identity.

### Both hub and non-hub enhancers are associated with active chromatin marks and master regulators

To further investigate molecular differences between hub and non-hub enhancers within hierarchical SEs, we compared the spatial patterns of histone marks among three enhancer groups: hub, non-hub and REs. Compared with non-hub enhancers, hub enhancers display no significant difference in H3K4me1 ChIP-seq signal (Figure 2A and Figure 2-figure supplement 1A), but are slightly more enriched for H3K27ac and DNase I hypersensitivity (Figure 2B,C and Figure 2-figure supplement 1B,C).

**Figure 2.**
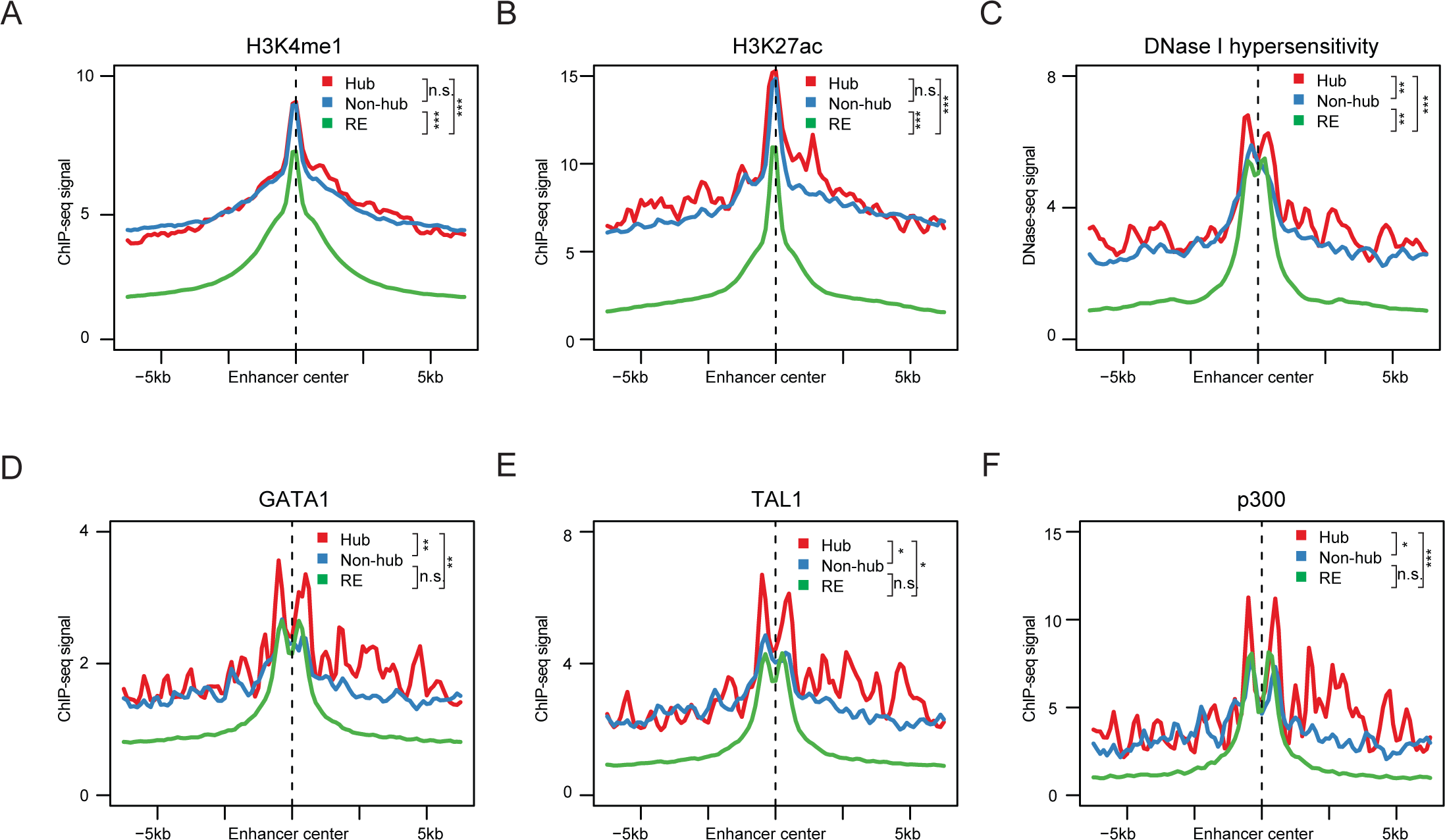
Chromatin landscapes at hub enhancers. (**A-F**) Spatial distribution of chromatin marks centered by enhancers in three groups in K562 cells, H3K4me1 (**A**), H3K27ac (**B**), DNase I hypersensitivity (**C**), master regulators GATA1 (**D**) and TAL1 (**E**), coactivator p300 (**F**). *P* values are calculated using Student’s t-test based on the ChIP-seq signal intensity within 1kb window centered by enhancers. \**P* < 0.05; \*\**P* < 0.01; \*\*\**P* < 0.001, n.s. not significant.

One of the hallmark features of SEs is the enrichment of cell type-specific master regulators and coactivators (Whyte et al., 2013). We then compared the distribution of transcription factor binding profiles. Hub enhancers contain moderate but significantly higher ChIP-seq signals for the binding of lineage-regulating master regulators than non-hub enhancers, such as GATA1 and TAL1 in K562 cells, and PAX5 and EBF1 in GM12878 cells (Figure 2D,E and Figure 2-figure supplement 1D,E). Hub enhancers also display increased occupancy of histone acetyltransferase p300, a coactivator associated with active enhancers (Figure 2F and Figure 2-figure supplement 1F). Taken together, these results demonstrate that hub and non-hub enhancers are characterized by quantitative differences in the occupancy of active enhancer-associated histone modifications and lineage-specifying transcription factors (TFs).

### Hub enhancers are distinctively enriched with cohesin and CTCF binding

Since hub and non-hub enhancers are defined based on the frequency of chromatin interactions, we next compared the occupancy of cohesin and CTCF, two factors essential for mediating long-range enhancer-promoter interactions and DNA looping (Ing-Simmons et al., 2015). To this end, we compared the enhancer groups with the ChIP-seq profiles for CTCF and two cohesin components, SMC3 and RAD21. Compared with non-hub enhancers, the occupancy of all three factors is markedly elevated at hub enhancers (Figure 3A-C and Figure 3-figure supplement 1A-C), consistent with a critical role of CTCF and cohesin in mediating chromatin interactions associated with hub enhancers. Importantly, while the role of CTCF in mediating chromatin organization, such as TADs, has been well established (Dixon et al., 2012), its association with SE constituents has not been previously reported. In fact, only a small fraction (6% in K562; 24% in GM12878) of hub enhancers overlap with known TAD boundaries (Figure 3D and Figure 3-figure supplement 1D), which is comparable to the genome-wide frequency of CTCF peaks overlapping with TAD boundaries, suggesting a TAD-independent role of CTCF.

**Figure 3.**
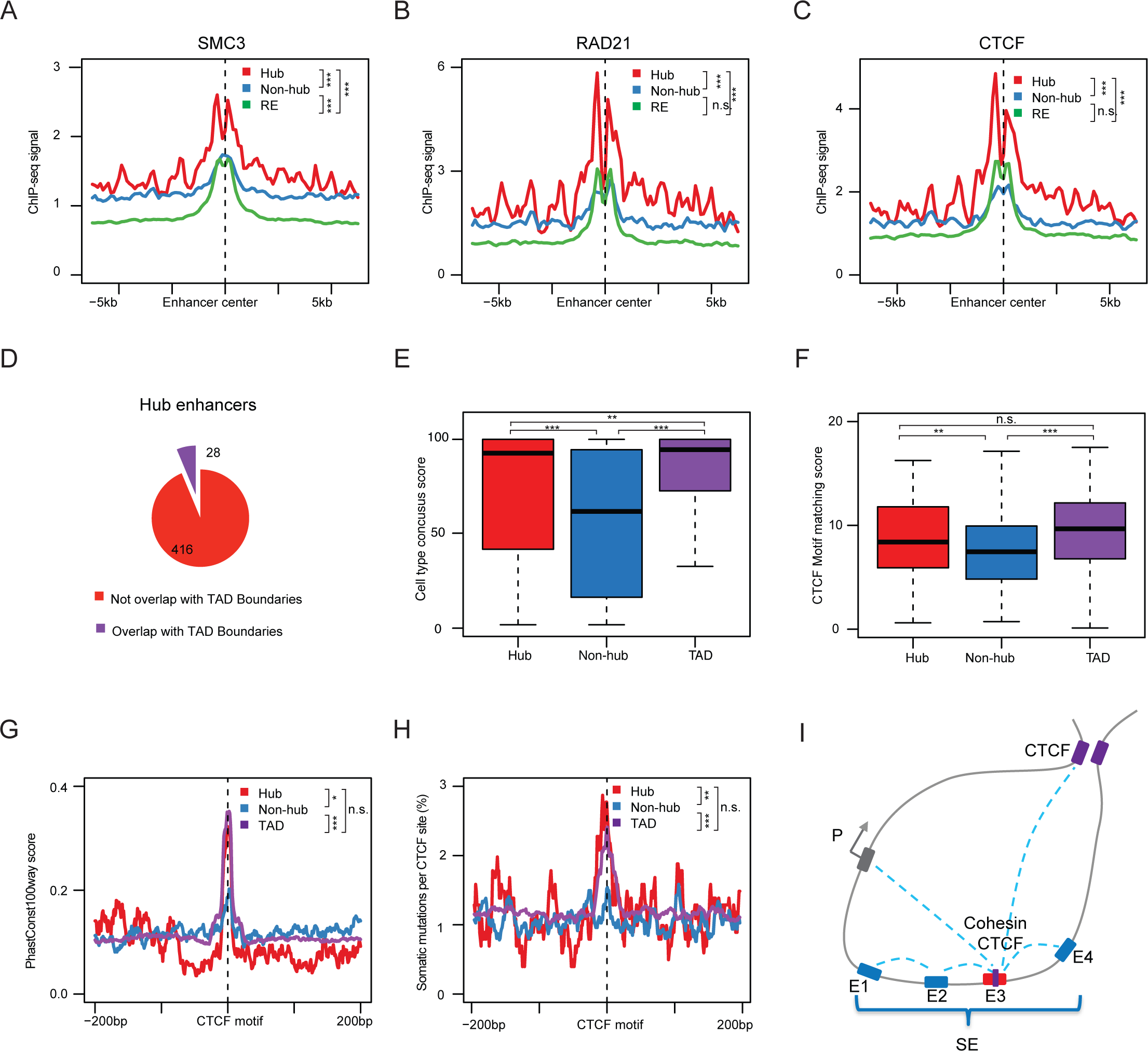
CTCF binding at hub enhancers within SEs hierarchy. (**A**-**C**) Spatial distribution of two cohesin components SMC3 (**A**) and RAD21(**B**), and CTCF (**C**), centered by enhancers in three groups. *P* values are calculated using Student’s t-test based on the ChIP-seq signal intensity of 1kb window centered by enhancers. \**P* < 0.05; \*\**P* < 0.01; \*\*\**P* < 0.001, n.s. not significant. (**D**) Percentage of hub enhancers with (purple) or without (red) overlapping with TAD boundaries collected from(Rao et al., 2014). The CTCF ChIP-seq peaks/motif-sites associated with hub enhancers overlapping with TAD boundaries were excluded for analysis in (**E-H**). (**E**) CTCF binding consensus across cell types in different contexts: hub (red), non-hub enhancers (blue) and TAD boundaries (purple). For each CTCF peak in K562, the consensus score (y-axis) was quantified as the percentage of cell types containing the same CTCF peak. *P* values are calculated using Student’s t test. \**P* < 0.05; \*\**P* < 0.01; \*\*\**P* < 0.001, n.s. not significant. (**F**) CTCF-motif-matching score (y-axis) of CTCF peaks. *P* values are calculated using Student’s t-test. \**P* < 0.05; \*\**P* < 0.01; \*\*\**P* < 0.001, n.s. *not significant.* (**G**) Sequence conservation around CTCF motif sites. The sitepro plots were centered by CTCF motif sites. *P* values are calculated using Student’s t-test based on the PhastConst100way score (y-axis) within CTCF motif sites. \**P* < 0.05; \*\**P* < 0.01; \*\*\**P* < 0.001, n.s. not significant. (**H**) Somatic mutation rate in cancers collected from IGGC around CTCF motif sites. The sitepro plots were centered by CTCF motif sites with 10bp smoothing window. *P* values are calculated using Fisher’s exact test based on overlap between CTCF motif sites and somatic mutation sites. \**P* < 0.05; \*\**P* < 0.01; \*\*\**P* < 0.001, n.s. not significant. (**I**) Model of the hierarchical organization of SEs containing both hub and non-hub enhancers. Hub enhancers are highly enriched with CTCF and cohesin binding, and functions as an organization hub to coordinate the non-hub enhancers and other distal regulatory elements within and beyond the SE.

To identify potential contextual differences between CTCF binding associated with distinct functions, we divided the CTCF ChIP-seq peaks into three non-overlapping subsets that overlap with hub enhancers, non-hub enhancers or TAD boundaries, respectively. To further distinguish CTCF binding at distinct regulatory regions, we excluded peaks that overlap with both hub enhancers and TAD boundaries (Figure 3d and Figure 3-figure supplement 1D). We first examined the cross cell-type variability of CTCF binding based on CTCF ChIP-seq signals in 55 cell types from ENCODE (T. E. P. Consortium, 2012). Consistent with previous studies (Dixon et al., 2012; Pope et al., 2014), we found that CTCF binding sites associated with TAD boundaries are highly conserved (Figure 3E and Figure 3-figure supplement 1E). In addition, within SEs, CTCF sites associated with hub enhancers are more conserved than those associated with non-hub enhancers. We hypothesized that the cell-type variability of CTCF binding may reflect the binding affinity of CTCF to its cognate sequences, which can be quantified by the motif-matching scores. Therefore, we compared the distribution of motif scores associated with different subsets of CTCF binding sites. The motif scores for CTCF sites associated with TAD boundaries and hub enhancers are higher than non-hub enhancer-associated CTCF sites, consistent with the CTCF ChIP-seq signal intensity (Figure 3F and Figure 3-figure supplement 1F). Of note, a similar pattern is observed for the genomic sequence conservation of CTCF binding sites as quantified by the phastCons100way score (Figure 3G and Figure 3-figure supplement 1G), suggesting that the cell-type variation associated with CTCF binding may be under evolutionary pressure.

Somatic mutations of TAD or insulated neighborhood boundaries have been reported in cancer (Flavahan et al., 2016; Hnisz, Weintraub, et al., 2016; Katainen et al., 2015). Consistently, we observed high frequency of somatic mutations in TAD boundary-associated CTCF sites using somatic mutations in different cancers from the ICGC database (International Cancer Genome et al., 2010). Hub-enhancer-associated CTCF sites display comparable rates of somatic mutations with TAD boundaries-associated CTCF sites, which are significantly higher than non-hub enhancer-associated CTCF sites (*P* = 9.0E-3 in K562, *P* = 2.3E-2 in GM12878, Figure 3G and Figure 3-figure supplement 1G). Our results suggest that genetic alterations of hub enhancer-associated CTCF sites may confer similar consequences as perturbations of TAD boundary-associated CTCF sites, such as activation of proto-oncogenes (Flavahan et al., 2016; Hnisz, Weintraub, et al., 2016). Taken together, our results support a model that hub enhancers have two molecularly and functionally related roles in SE hierarchy (Figure 3I). Hub enhancers act as ‘conventional’ enhancers to activate gene expression through the recruitment of lineage-specifying transcriptional regulators and coactivators. In addition, they act as ‘organizational’ hubs to mediate and/or facilitate long-range chromatin interactions through the recruitment of cohesin and CTCF complexes.

### Hub enhancers are enriched for genetic variants associated with cell-type-specific gene expression and diseases

Genetic variations colocalized with regulatory genomic elements often associate with variation in expression of the linked target genes. As such, expression quantitative trait loci (eQTL) enrichment analysis serves as an objective and quantitative metric to evaluate regulatory potential. We compared the frequencies of eQTLs that are significantly associated with gene expression from the GTEx eQTL database (G. T. Consortium, 2013) with hub, non-hub and regular enhancers (Figure 4A and Figure 4- figure supplement 1A). We observed that SEs are more enriched with eQTLs than regular enhancers (Figure 4-figure supplement 2A). Importantly, within SEs, hub enhancers are more enriched with eQTLs compared to non-hub enhancers (Figure 4A and Figure 4-figure supplement 1A). The difference is more apparent in the comparison using eQTLs identified in blood cells (Figure 4B,C and Figure 4-figure supplement 1B,C).

**Figure 4.**
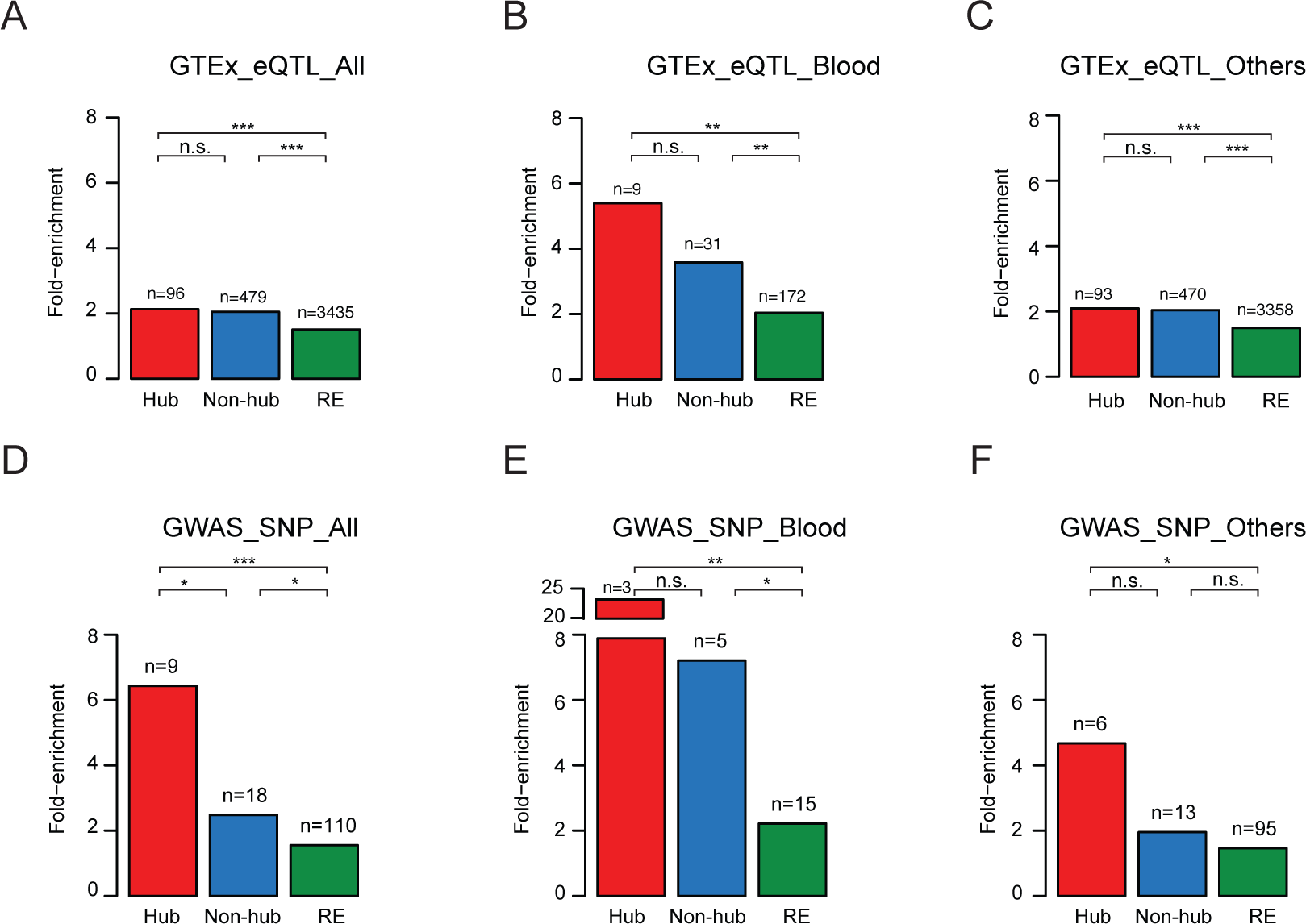
Enrichment of genetic variants associated with cell-type-specific gene expression and diseases in hub enhancers. (**A-C**) Enrichment of the eQTLs curated in GTEx in the enhancers in three groups in K562 cells, using randomly selected genome regions as control (see Methods). The GTEx eQTL identified in all tissues (**A**) were separated into two subsets, identified in whole blood (**B**) or other tissues (**C**). The number of enhancers overlap in each group with eQTLs were labelled on each bar. *P* values are calculated using Fisher’s exact test. \**P* < 0.05; \*\**P* < 0.01; \*\*\**P* < 0.001, n.s. not significant. (**D-F**) Enrichment of the disease or traits-associated SNPs curated in GWAS catalog in the enhancers in three groups in K562 cells, using randomly selected genome regions as control (see Methods). The GWAS SNPs associated all diseases/traits (**D**), were separated into two subsets, associated with blood-related diseases/traits (**E**) or other traits (**F**). The number of enhancers overlap in each group with SNPs were labelled on each bar. *P* values are calculated using Fisher’s exact test. \**P* < 0.05; \*\**P* < 0.01; \*\*\**P* < 0.001, n.s. not significant.

To gain insights into the function of hub enhancers, we next compared the enhancer groups with genome-wide association study (GWAS)-identified disease-associated genetic variants. Specifically, we analyzed the enrichment of single-nucleotide polymorphisms (SNPs) linked to diverse phenotypic traits and diseases in the GWAS catalog (Welter et al., 2014). Whereas REs are 1.6- and 1.9-fold more enriched with GWAS SNPs relative to genome background in K562 and GM12878 cells, respectively, the enrichment scores for SEs are significantly higher (2.7- and 4.8-fold, respectively) (Figure 4-figure supplement 2A). The enrichment of GWAS SNPs at SEs is consistent with previous studies that SEs are enriched with disease-associated variants (Hnisz et al., 2013; Maurano et al., 2012). Importantly, within SEs, hub enhancers display significantly higher enrichment (6.4- and 6.8-fold) than non-hub enhancers (2.5- and 4.5-fold) or REs (Figure 4D and Figure 4-figure supplement 1D). Furthermore, hub enhancers in K562 cells display the highest enrichment of GWAS SNPs associated with blood traits (22.4-fold, Figure 4E,F), indicating that hub enhancers enrich for cell-type-specific diseases-associated variants. We also found the hub enhancers defined by different thresholds of H-scores display similar enrichment of eQTLs and GWAS SNPs (Figure 4-figure supplement 2B,C), indicating that the properties of hub enhancers are not dependent on the specific threshold of H-score. Taken together, our studies demonstrate that hub enhancers within SEs are most significantly enriched with genetic variants associated with diseases and cell-type-specific gene expression, supporting their roles in the control of cell identity and disease.

To test the robustness of our method, we repeated our analysis to define hierarchical SEs and hub enhancers based on CTCF-mediated ChIA-PET datasets in K562 and GM12878 cells (Tang et al., 2015) (see Methods). We observed that 102 of 188 hierarchical SEs in K562 and 227 of 427 hierarchical SEs in GM12878 defined by ChIA-PET datasets overlap with those defined by Hi-C data (*P* < 2.2E-16 in both K562 and GM12878, Fisher’s exact test, Figure 4-figure supplement 3A). The hub enhancers within the hierarchical SEs shared by both data types also significantly overlap (*P* <2.2E-16 in both K562 and GM12878, Fisher’s exact test). Similar to previous analysis, we observed that hub enhancers defined by ChIA-PET data were also more enriched with disease-associated variants compared to non-hub enhancers (Figure 4-figure supplement 3B). The consistency between analyzing two independent experimental platforms (Hi-C and ChIA-PET), as well as between analyzing two distinct cell types (K562 and GM12878), strongly indicates that our approach is robust and generally applicable.

### *In situ* genome editing reveals distinct requirement of hub vs non-hub enhancers in SE function

Since the structural organization of chromatin plays a critical role in establishing enhancer activities, we then compared the regulatory potential of hub and non-hub enhancers subjected to genetic perturbation. In prior work, we applied CRISPR/Cas9 based genome-editing to systematically dissect the functional hierarchy of an erythroid-specific SE controlling the *SLC25A37* gene encoding the mitochondrial transporter critical for iron metabolism (Huang et al., 2016). Following deletion of each of the three constituent enhancers alone or in combination, we identified a functionally ‘dominant’ enhancer responsible for the vast majority of enhancer activity (Huang et al., 2016). Of note, we found that this ‘dominant’ enhancer is identified as a hub enhancer and associated with significantly higher chromatin interactions compared to the neighboring non-hub enhancers (Figure 5-figure supplement 1A). These studies provide initial evidence that hub enhancers may be more transcriptionally potent than non-hub enhancers in gene activation.

To further establish the functional roles of hub enhancers, we performed experimental validation of hierarchical SEs identified in K562 cells based on the predictions of our model. We first employed CRISPR interference (CRISPRi) in which the nuclease-dead Cas9 protein (dCas9) is fused to a KRAB (Kruppel-associated box) transcriptional repressor domain (Gilbert et al., 2014; Thakore et al., 2015; Xie, Duan, Li, Zhou, & Hon, 2017). Upon co-expression of sequence-specific single guide RNAs (sgRNAs) targeting individual hub or non-hub enhancers in K562 cells, we measured the expression of SE-linked target genes as a readout for the functional requirement for SE activity. We focused on two representative SE clusters located in the proximity of the *MYO1D* and *SMYD3* genes (Figure 5-figure supplement 1B,C and Figure 5A,B). Both SEs were predicted to contain hierarchical structure (H-score=2.2 and 1.6 respectively), while their nearest target genes *MYO1D* and *SMYD3* are highly expressed in K562 cells. Moreover, both SEs contain hub and non-hub enhancers within a defined TAD domain (Figure 5-figure supplement 1B,C). Importantly, whereas CRISPRi-mediated repression of the two non-hub enhancers at the *MYO1D* SE led to modest downregulation (3.1-fold) of *MYO1D* expression, repression of the hub enhancer significantly decreased *MYO1D* expression by 8.3-fold (Figure 5C,D). Similarly, CRISPRi-mediated repression of the hub enhancer located in the *SMYD3* SE cluster resulted in more profound downregulation of *SMYD3* expression compared to the non-hub enhancer (Figure 5E).

**Figure 5.**
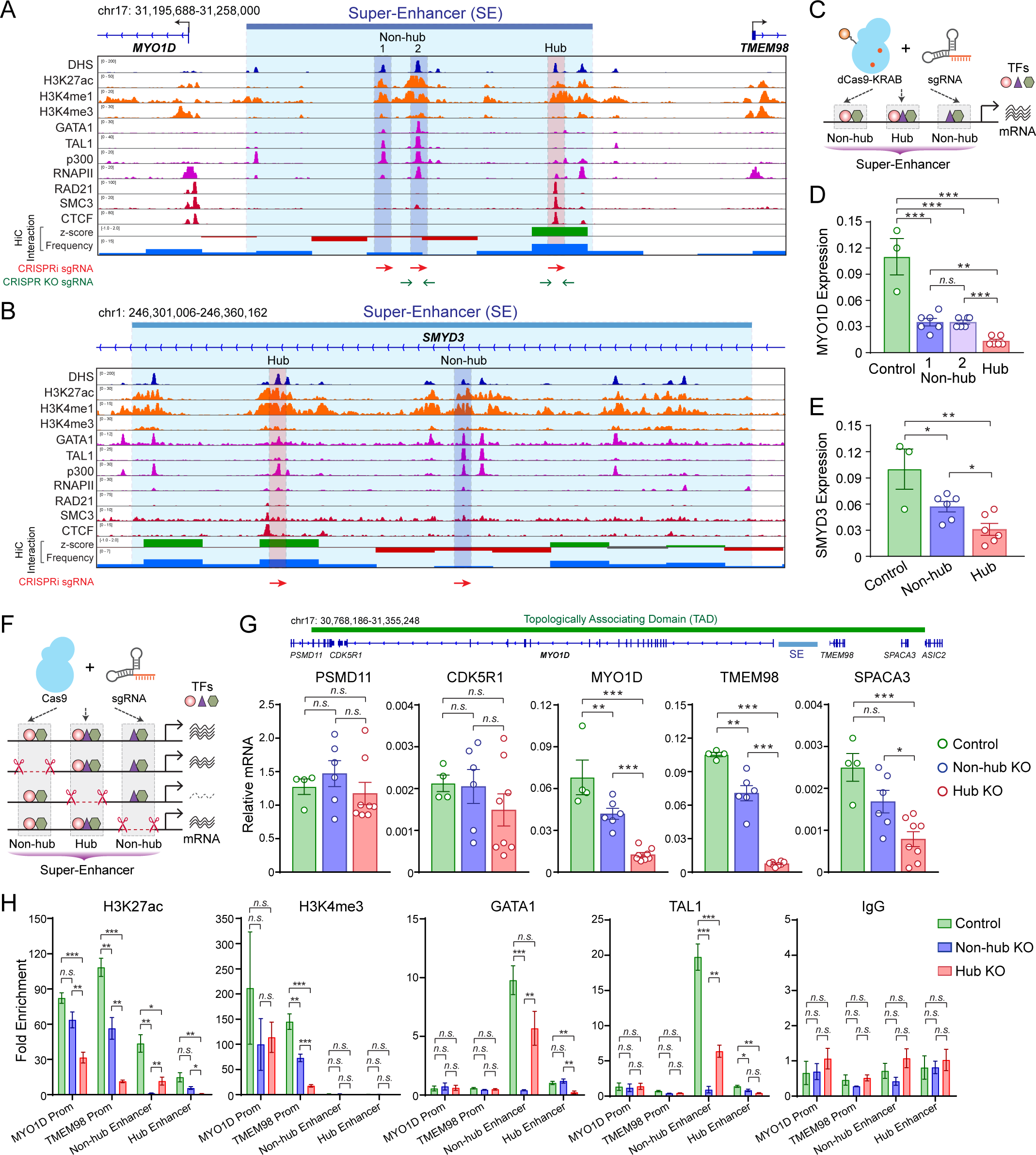
*In situ* genome editing reveals distinct requirement of hub vs non-hub enhancers in SE function. (**A**) Chromatin signatures and TF occupancy at the MYO1D SE locus in K562 cells are shown. The identified hub and non-hub enhancers are depicted by red (hub) and blue (non-hub) lines, respectively. The Hi-C chromatin interaction z-score and frequency at 5kb resolution is shown at the bottom (see Methods). The positions of sgRNAs used for CRISPRi or CRISPR/Cas9-mediated knockout analyses are shown as arrowheads. (**B**) Chromatin signatures and TF occupancy at the SMYD3 SE locus in K562 cells are shown. (**C**) Schematic of CRISPRi-mediated repression of hub or non-hub enhancers. (**D,E**) Expression of MYO1D and SMYD3 mRNA in untreated (control), CRISPRi-mediated repression of hub or non-hub enhancers. The mRNA expression levels related to GAPDH are shown. Each colored circle represents an independent biological replicate experiment. Results are means ± SEM. *P* values are calculated by two-sided Student’s t-test. \**P* < 0.05, \*\**P* < 0.01, \*\*\**P* < 0.001, n.s. not significant. (**F**) Schematic of CRISPR/Cas9-mediated knockout of hub or non-hub enhancers. (**G**) Expression of all genes within the SE-containing TAD domain in unmodified (control), CRISPR/Cas9-mediated knockout of hub or non-hub enhancers. The mRNA expression levels relative to GAPDH are shown. Each colored circle represents an independent single-cell-derived biallelic enhancer knockout clone. A schematic of the SE-containing TAD domain and associated genes are shown on the top. Results are means ± SEM. *P* values are calculated by a two-sided Student’s t-test. \**P* < 0.05, \*\**P* < 0.01, \*\*\**P* < 0.001, n.s. not significant. (**H**) ChIP-qPCR analysis of H3K27ac, H3K4me3, GATA1, TAL1 and IgG (negative control) in unmodified (control), hub or non-hub enhancer knockout cells. Primers against MYO1D and TMEM98 promoters, hub and non-hub enhancers, and a negative control genome region (chr2:211,337,339-211,337,429) are used. The results are shown as fold enrichment of the ChIP signals against the negative control region as means ± SEM of four independent experiments. *P* values are calculated by a two-sided Student’s t-test. \**P* < 0.05, \*\**P* < 0.01, \*\*\**P* < 0.001, n.s. not significant.

To further interrogate the role of hub versus non-hub enhancers in SE structure and function *in situ*, we employed CRISPR/Cas9-mediated genome engineering to delete individual hub or non-hub enhancers with paired sgRNAs flanking the enhancer elements at the *MYO1D* SE (Figure 5F). We observed that 3 of 5 genes within the SE-containing TAD domain (*MYO1D*, *TMEM98* and *SPACA3*) displayed significant downregulation in mRNA expression, whereas the other two genes (*PSMD11* and *CDK5R1*) remained unaffected (Figure 5G and Figure 5-figure supplement 1B), suggesting that the *MYO1D* SE regulates only a subset of genes within the same TAD domain. Furthermore, knockout of the hub enhancer resulted in more significant downregulation (5.4, 14.0 and 3.2-fold related to control; *P* < 0.001) of *MYO1D*, *TMEM98* and *SPACA3* genes compared to the non-hub enhancers (1.6, 1.5 and 1.5- fold), respectively, consistent with a prominent role of hub enhancers in SE activity. To measure the effects on the local chromatin landscape, we performed ChIP experiments in control, hub and non-hub enhancer knockout cells (Figure 5H). We observed that knockout of the non-hub enhancer had only a subtle effect on the enhancer-associated histone mark (H3K27ac) and binding of master TFs (GATA1 and TAL1) at the promoter or enhancer regions of SE-linked *MYO1D* and *TMEM98* genes. In contrast, knockout of the hub enhancer led to marked downregulation, or near absence, of H3K27ac, H3K4me3 and GATA1/TAL1 binding at neighboring enhancers or promoters. These results demonstrate that hub enhancers are functionally more potent than neighboring non-hub enhancers in directing transcriptional activation of SE-linked gene targets.

Taken together, our *in situ* genome editing analysis of multiple representative SE clusters provides compelling evidence that at least a subset of SEs are composed of a hierarchical structure containing hub and non-hub enhancer elements, whereby hub enhancers are functionally indispensable for SE activities.

## Discussion

SE assignment provides a means to identify regulatory regions near important genes that regulate cell fate (Pott & Lieb, 2015). However, it has remained unclear how SEs function and the extent to which they are distinct from more conventional enhancers. As such, the challenge has been to ascribe functional features uniquely associated with SEs, and account for how the activities of the constituent elements are coordinated for SE function (Pott & Lieb, 2015). Here, we have developed a systematic approach to interrogate the structural hierarchy of SE constituent elements. First, we observed that only a subset of SEs contains a hierarchical structure, which is consistent with previous findings that SEs are intrinsically heterogeneous, with a large fraction of SEs containing 3 or fewer constituent elements (Pott & Lieb, 2015). Such heterogeneity may provide one explanation for an apparent paradox in the literature (Dukler, Gulko, Huang, & Siepel, 2016; Pott & Lieb, 2015). For example, recent studies by our group and others provided evidence that SEs may be composed of a hierarchy of enhancer constituents that coordinately regulate gene expression (Canver et al., 2015; Fulco et al., 2016; Hnisz et al., 2015; Huang et al., 2016; H. Y. Shin et al., 2016). On the other hand, other examples suggest that some SEs may not contain hierarchical structures and the SE constituents contribute additively to gene activation (Hay et al., 2016; Moorthy et al., 2017). Within hierarchical SEs, we identified those hub enhancers associated with an unusually high frequency of long-range chromatin interactions, suggesting that these elements may play an important role in maintaining the structure of SEs. Moreover, hub enhancers are significantly more enriched with eQTL and GWAS-identified genetic variations, and functionally more potent for gene activation than neighboring non-hub enhancers within the same SEs. Hence, our results support a model in which the structural hierarchy of SEs is predictive of functional hierarchy.

We observed that CTCF binding is highly enriched at hub enhancers compared to other constituent elements. CTCF has an established role in orchestrating genome structure (Phillips & Corces, 2009). The prevailing model posits that the primary function of CTCF is to maintain the boundaries of topological domains and the insulated neighborhoods (Hnisz, Day, & Young, 2016). Beyond this, our results suggest that CTCF plays additional, yet important, roles in organizing the structural hierarchy of SEs. We speculate that hierarchical organization may be established in a step-wise manner during development through coordinated interactions between CTCF and cell-type specific regulators. Disruption of the hierarchical organization of SE structures may impair SE function and predispose to pathological conditions (Flavahan et al., 2016; Hnisz, Weintraub, et al., 2016; Katainen et al., 2015). Consistent with this model, we found that hub-enhancer-associated CTCF sites display a significantly higher frequency of somatic mutation than non-hub enhancer-associated CTCF sites. Thus, it will be important to investigate chromatin interaction landscapes at both single gene and genomic levels in cancer cells harboring somatic mutations in CTCF sites.

At present, Hi-C or ChIA-PET datasets are limited in resolution and available cell types, which presents a significant challenge for further investigation of structural organization within SEs across cell types and cellular conditions. However, the recent development of new technologies, including Hi-ChIP, GAM and capture Hi-C (Beagrie et al., 2017; Mumbach et al., 2016; Schoenfelder et al., 2015), promises to enhance the quality and efficiency of data collection for 3D genome structures in various cell types. At the same time, improved methods for functional validation are also being rapidly developed, such as high-resolution CRISPR/Cas9 mutagenesis (Canver et al., 2017; Canver et al., 2015; Diao et al., 2017). With anticipated availability of additional chromatin interaction datasets, the computational method we describe here should find wide applications to the systematic investigation of the functional and structural organization of regulatory elements, including and beyond SEs. Findings from these studies will provide mechanistic insights into the genetic and epigenetic components of human genome in development and disease.

## Materials and Methods

### Identification of SEs

ChIP-seq data of H3K27ac in K562 and GM12878 cells were downloaded from ENCODE (T. E. P. Consortium, 2012). All data were in the human genome version hg19. MACS2 (Zhang et al., 2008) was used to identify H3K27ac peaks with a threshold Q-value=1.0E-5. H3K27ac peaks were used to define the enhancer boundary, followed by further filtering based on the criteria: (1) excluding H3K27ac peaks that overlapped with ENCODE blacklisted genomic regions (T. E. P. Consortium, 2012); and (2) excluding H3K27ac peaks that were located within +/-2kb region of any Refseq annotated gene promoter. The remaining H3K27ac peaks were defined as enhancers. Then, SEs were identified by using the ROSE (Rank Ordering of Super-Enhancers) algorithm (Loven et al., 2013; Whyte et al., 2013) based on the H3K27ac ChIP-seq signal with the default parameters.

### Analysis of Hi-C data

The 5kb resolution intra-chromosomal raw interaction matrix in K562 and GM12878 cells were downloaded from a public dataset (Rao et al., 2014). The statistically significant chromatin interactions were detected as previous (Huang et al., 2015). Briefly, the raw interaction matrix was normalized by using the ICE algorithm (Imakaev et al., 2012), as implemented in the Hi-Corrector package (Li, Gong, Li, Alber, & Zhou, 2015), to remove biases (Imakaev et al., 2012; Peng et al., 2013). Fit-Hi-C (Ay, Bailey, & Noble, 2014) was used to identify statistically significant intra-chromosomal interactions, using the parameter setting ‘-U=2000000, -L=10000’ along with the threshold of FDR=0.01. The interaction frequency for each 5kb bin was calculated as the number of significant chromatin interactions associated with the bin. The list of TADs in K562 and GM12878 cells were downloaded from the supplementary data (Rao et al., 2014).

### Analysis of chromatin mark distributions

ChIP-seq of histone marks (H3K27ac, H3K4me1) and transcription factors/co-activators (GATA1, TAL1, PAX5, EBF1, p300, CTCF, SMC3, RAD21), DNase-seq in K562, GM12878 cells were downloaded from ENCODE (T. E. P. Consortium, 2012). Replicate data were merged if available. The sitepro plots for chromatin marks were plotted based on the binned density matrix range from +/-5kb centered by enhancer generated by using the CEAS software (H. Shin, Liu, Manrai, & Liu, 2009).

### Analysis of CTCF related datasets

Genome-wide CTCF peak locations in 55 cell types, including K562 and GM12878 cells, were downloaded from ENCODE (T. E. P. Consortium, 2012). For each CTCF peak in K562 or GM12878, the cell type consensus score was defined as the percentage of cell types in which the peak was detected.

CTCF motif information, represented as a position weight matrix, was downloaded from the JASPAR database (Mathelier et al., 2014). For each CTCF peak in K562 or GM12878, the corresponding maximum motif-matching score was evaluated by using the HOMER software (Heinz et al., 2010).

The phastCons scores (Siepel et al., 2005) for multiple alignments of 99 vertebrate genomes to the human genome were downloaded from the UCSC Genome Browser. The sitepro plots of conservation score were plotted within +/-200bp centered by CTCF motif sites.

Known somatic mutation loci in cancer were downloaded from International Cancer Genome Consortium (ICGC) (International Cancer Genome et al., 2010) Data Portal under release 23. The sitepro plots of mutation frequencies were plotted within +/- 200bp centered by CTCF motif sites with a 10bp smoothing window.

### Enrichment analysis of GWAS SNPs and eQTLs

The SNPs curated in GWAS Catalog (Welter et al., 2014) were downloaded through the UCSC Table Browser (Karolchik et al., 2004). The subset of blood-associated GWAS SNPs was selected as those associated with at least one of the following keywords in the “trait” field: ‘Erythrocyte’, ‘F-cell’, ‘HbA2’, ‘Hematocrit’, ‘Hematological’, ‘Hematology’, ‘Hemoglobin’, ‘Platelet’, ‘Blood’, ‘Anemia’, ‘sickle cell disease’, ‘Thalassemia’, ‘Leukemia’, ‘Lymphoma’, ‘Lymphocyte’, ‘B cell ‘, ‘B-cell’, ‘Lymphoma’, ‘Lymphocyte’, and ‘White blood cell’. Enrichment analysis was carried out as described previously (Huang et al., 2015), using random permutation as control.

Statistically significant eQTL loci in multiple tissues were downloaded from the Genotype-Tissue Expression (GTEx) database (Accession phs000424.v6.p1) (G. T. Consortium, 2013). Blood-associated eQTLs were those identified in the whole blood.

### Analysis of ChIA-PET dataset

CTCF-mediated ChIA-PET data were downloaded from ENCODE (for K562) and from the publication website (Tang et al., 2015) (for GM12878), respectively. The interaction frequency of each 5kb bin was calculated as the number of chromatin interactions associated the PET clusters located in the bin.

### Data visualization

The ChIP-seq signal and peaks were visualized using Integrative Genomics Viewer (IGV) (Robinson et al., 2011).

### Cell culture

K562 cells were obtained from the American Tissue Collection Center (ATCC). K562 cells were cultured in RPMI1640 medium supplemented with 10% FBS and 1% penicillin-streptomycin.

### CRISPR/Cas9-Mediated Interference (CRISPRi) of enhancer elements

The CRISPR interference (CRISPRi) system was used to investigate the function of enhancer elements following published protocol with modifications (Gilbert et al., 2014; Thakore et al., 2015). Briefly, sequence-specific sgRNAs for site-specific interference of genomic targets were designed following described guidelines, and sequences were selected to minimize off-target effect based on publicly available filtering tools (http://crispr.mit.edu/). Oligonucleotides were annealed in the following reaction: 10 μM guide sequence oligo, 10 μM reverse complement oligo, T4 ligation buffer (1X), and 5U of T4 polynucleotide kinase (New England Biolabs) with the cycling parameters of 37°C for 30 min; 95°C for 5 min and then ramp down to 25°C at 5°C/min. The annealed oligos were cloned into pLV-hU6-sgRNA-hUbC-dCas9-KRAB-T2a-Puro vector (Addgene ID: 71236) using a Golden Gate Assembly strategy including: 100 ng of circular pLV plasmid, 0.2 μM annealed oligos, 2.1 buffer (1X) (New England Biolabs), 20 U of BsmBI restriction enzyme, 0.2 mM ATP, 0.1 mg/ml BSA, and 750 U of T4 DNA ligase (New England Biolabs) with the cycling parameters of 20 cycles of 37°C for 5 min, 20°C for 5 min; followed by 80°C incubation for 20 min. Then K562 cells were transduced with lentivirus to stably express dCas9-KRAB and sgRNA. To produce lentivirus, we plated K562 cells at a density of 3.0 × 106 per 10 cm plate in high-glucose DMEM supplemented with 10% FBS and 1% penicillin-streptomycin. The next day after seeding, cells were cotransfected with the appropriate dCas9-KRAB lentiviral expression plasmid, psPAX2 and pMD2.G by PEI (Polyethyleneimine). After 8 h, the transfection medium was replaced with 5 mL of fresh medium. Lentivirus was collected 48 h after the first media change. Residual K562 cells were cleared from the lentiviral supernatant by filtration through 0.45 μm cellulose acetate filters. To facilitate transduction, we added the PGE2 (Prostaglandin E2) to the viral media at a concentration of 5 μM. The day after transduction, the medium was changed to remove the virus, and 1 μg/ml puromycin was used to initiate selection for transduced cells. The positive cells were expanded and processed for gene expression analysis.

### CRISPR/Cas9-mediated knockout of enhancer elements

The CRISPR/Cas9 system was used to introduce deletion mutations of enhancer elements in K562 cells following published protocols (Canver et al., 2014; Cong et al., 2013; Mali et al., 2013). Briefly, the annealed oligos were cloned into pSpCas9(BB) (pX458; Addgene ID: 48138) vector using a Golden Gate Assembly strategy. To induce segmental deletions of candidate regulatory DNA regions, four CRISPR/Cas9 constructs were co-transfected into K562 cells by nucleofection using the ECM 830 Square Wave Electroporation System (Harvard Apparatus, Holliston, MA). Each construct was directed to flanking the target genomic regions. To enrich for deletion, the top 1-5% of GFP-positive cells were FACS sorted 48-72 h post-transfection and plated in 96-well plates. Single cell derived clones were isolated and screened for CRISPR-mediated deletion of target genomic sequences. PCR amplicons were subcloned and analyzed by Sanger DNA sequencing to confirm non-homologous end-joining (NHEJ)-mediated repair upon double-strand break (DSB) formation. The positive single-cell-derived clones containing the site-specific deletion of the targeted sequences were expanded for processed for gene expression analysis. The sequences of sgRNAs and genotyping PCR primers are listed in Figure 5-figure supplement 2.

### Chromatin immunoprecipitation (ChIP)

ChIP experiments were performed as described with modifications (Huang et al., 2016). Briefly, 2~5 x 10^6^ cells were crosslinked with 1% formaldehyde for 5 min at room temperature. Chromatin was sonicated to around 500 bp in RIPA buffer (10 mM Tris-HCl, 1 mM EDTA, 0.1% sodium deoxycholate, 0.1% SDS, 1% Triton X-100, 0.25% sarkosyl, pH 8.0) with 0.3 M NaCl. Sonicated chromatin were incubated with 2μg antibody at 4oC. After overnight incubation, protein A or G Dynabeads (Invitrogen) were added to the ChIP reactions and incubated for four additional hours at 4oC to collect the immunoprecipitated chromatin. Subsequently, Dynabeads were washed twice with 1 ml of RIPA buffer, twice with 1 ml of RIPA buffer with 0.3 M NaCl, twice with 1 ml of LiCl buffer (10 mM Tris-HCl, 1 mM EDTA, 0.5% sodium deoxycholate, 0.5% NP-40, 250 mM LiCl, pH 8.0), and twice with 1 ml of TE buffer (10 mM Tris-HCl, 1 mM EDTA, pH 8.0). The chromatin were eluted in SDS elution buffer (1% SDS, 10 mM EDTA, 50 mM Tris-HCl, pH 8.0) followed by reverse crosslinking at 65oC overnight. ChIP DNA were treated with RNaseA (5 μg/ml) and protease K (0.2 mg/ml), and purified using QIAquick Spin Columns (Qiagen). The purified ChIP DNA was quantified by real-time PCR using the iQ SYBR Green Supermix (Bio-Rad). The following antibodies were used: H3K27ac (ab4729, Abcam), H3K4me3 (04-745, Millipore), IgG (12-370, Millipore), GATA1 (ab11852, Abcam), TAL1 (sc-12984, Santa Cruz Biotechnology).

### Replicates

The biological replicates are defined as experiments performed using independently isolated biological samples grown/treated under the same conditions. The technical replicates are defined as experiments performed using the same sample (after all preparatory techniques) and analyzed in multiple times. For the CRISPR/Cas9-mediated knockout (KO) of hub or non-hub enhancers (Figure 5C-G), three or four independent single cell-derived homozygous KO clones were analyzed, each with two technical replicates. The unmodified control cells were analyzed as two independent biological replicate experiments, each with two technical replicates. For the ChIP-qPCR analysis (Figure 5H), the results are shown as means ± SEM of two biological replicates, each with two technical replicate measurements. All experimental data points including outliers were included in the data analysis.

## Authors’ contributions

J.H., J.X. and G.C.Y. conceived and designed the experiments. J.H. and Y.Z. performed bioinformatic analyses. K.L. and X.L. performed experimental validation. J.H., J.X., G.C.Y., K.L., W.C. and S.H.O. wrote the manuscript. J.X. and G.C.Y. supervised the project.

## Acknowledgements

We thank Drs. Shiqi Xie and Gary Hon for providing the dCas9-KRAB construct. This work was supported by NIH/NIDDK grants K01DK093543, R03DK101665 and R01DK111430, by a Cancer Prevention and Research Institute of Texas (CPRIT) New Investigator award (RR140025), by the American Cancer Society (IRG-02-196) award and the Harold C. Simmons Comprehensive Cancer Center at UT Southwestern, and by an American Society of Hematology Scholar Award (to J.X.). G.C.Y.’s research was supported by the NIH/NHLBI grant R01HL119099. We thank Dr. Alan Cantor and members of the Yuan Lab for helpful discussions.

## Supplementary Figures

**Figure 1-figure supplement 1.**
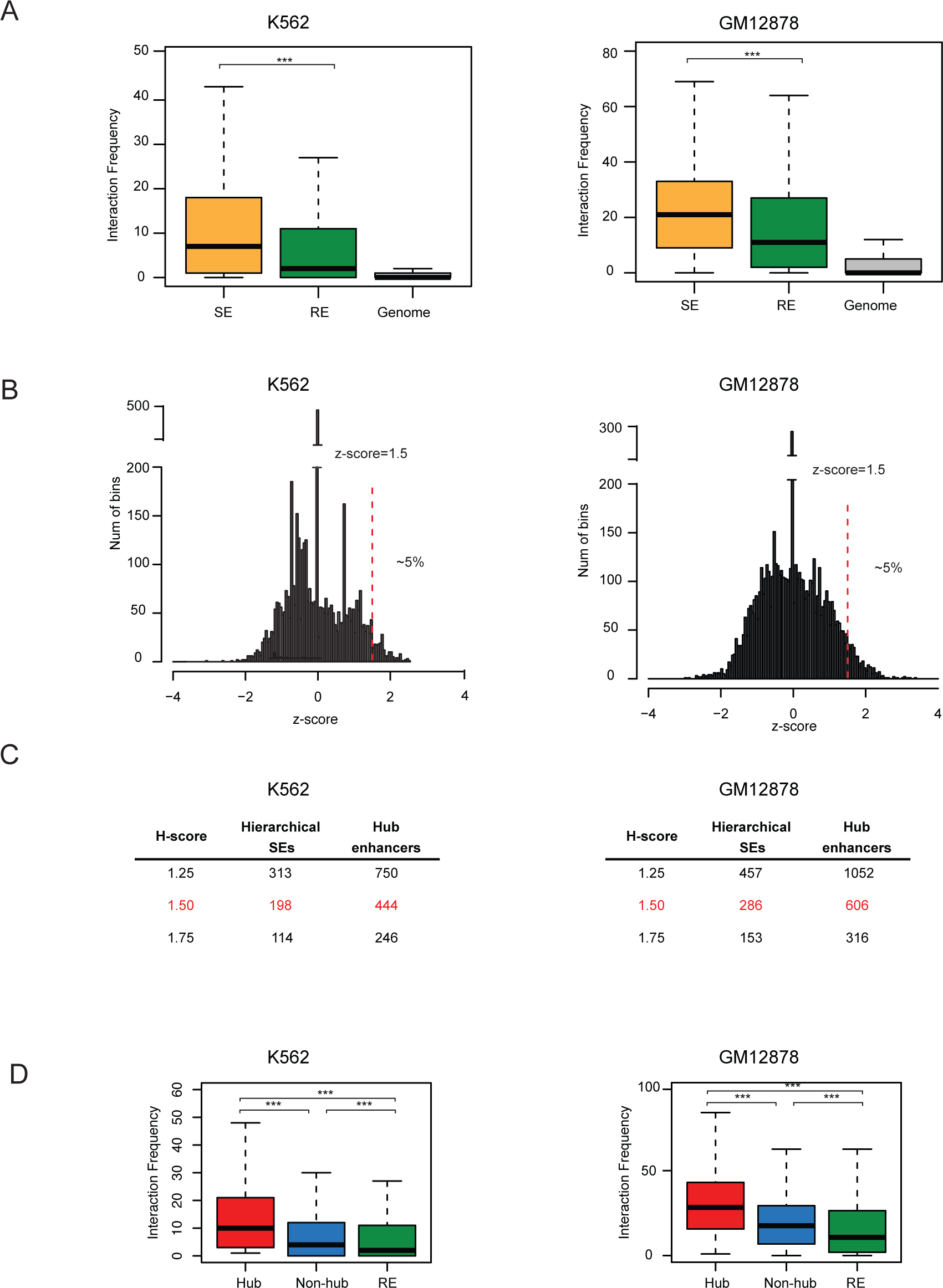
Definition of hierarchical SEs and hub enhancers using chromatin interactions in K562 and GM12878 cells. (**A**) Chromatin interactions frequency for 5kb bins overlapping with SEs (yellow), REs (green), using randomly selected genome 5kb bins as control (gray) in K562 and GM12878 cells. *P* values are calculated using Student’s t-test. \**P* < 0.05; \*\**P* < 0.01; \*\*\**P* < 0.001, n.s. not significant. (**B**) Distribution of z-score of 5kb bins in all SEs. The dashed line represents the threshold value of H-score = 1.5, which roughly corresponds to the 95^th^ percentile of z-scores. (**C**) Hierarchical SEs and hub enhancers defined using different thresholds of H-score. (**D**) The frequency of chromatin interaction of enhancers in three groups of enhancers (red for hub enhancers, blue for non-hub enhancers, green for regular enhancers). *P* values are calculated using Student’s t-test. \**P* < 0.05; \*\**P* < 0.01; \*\*\**P* < 0.001, n.s. not significant.

**Figure 1-figure supplement 2.**
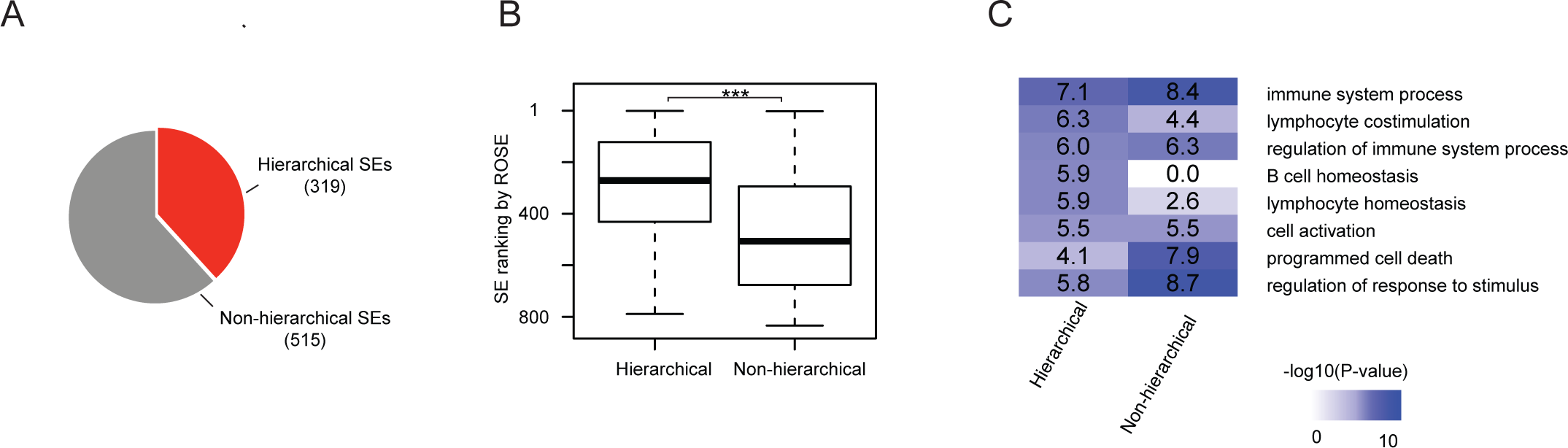
Hierarchical and non-hierarchical SEs in GM12878 cells. (**A**) Proportion of hierarchical and non-hierarchical SEs. (**B**) The ROSE ranking of hierarchical and non-hierarchical SEs. *P* value is calculated using Wilcoxon rank-sum test. \**P* < 0.05; \*\**P* < 0.01; \*\*\**P* < 0.001, n.s. not significant. (**C**) GREAT functional analysis of hierarchical and non-hierarchical SEs.

**Figure 2-figure supplement 1.**
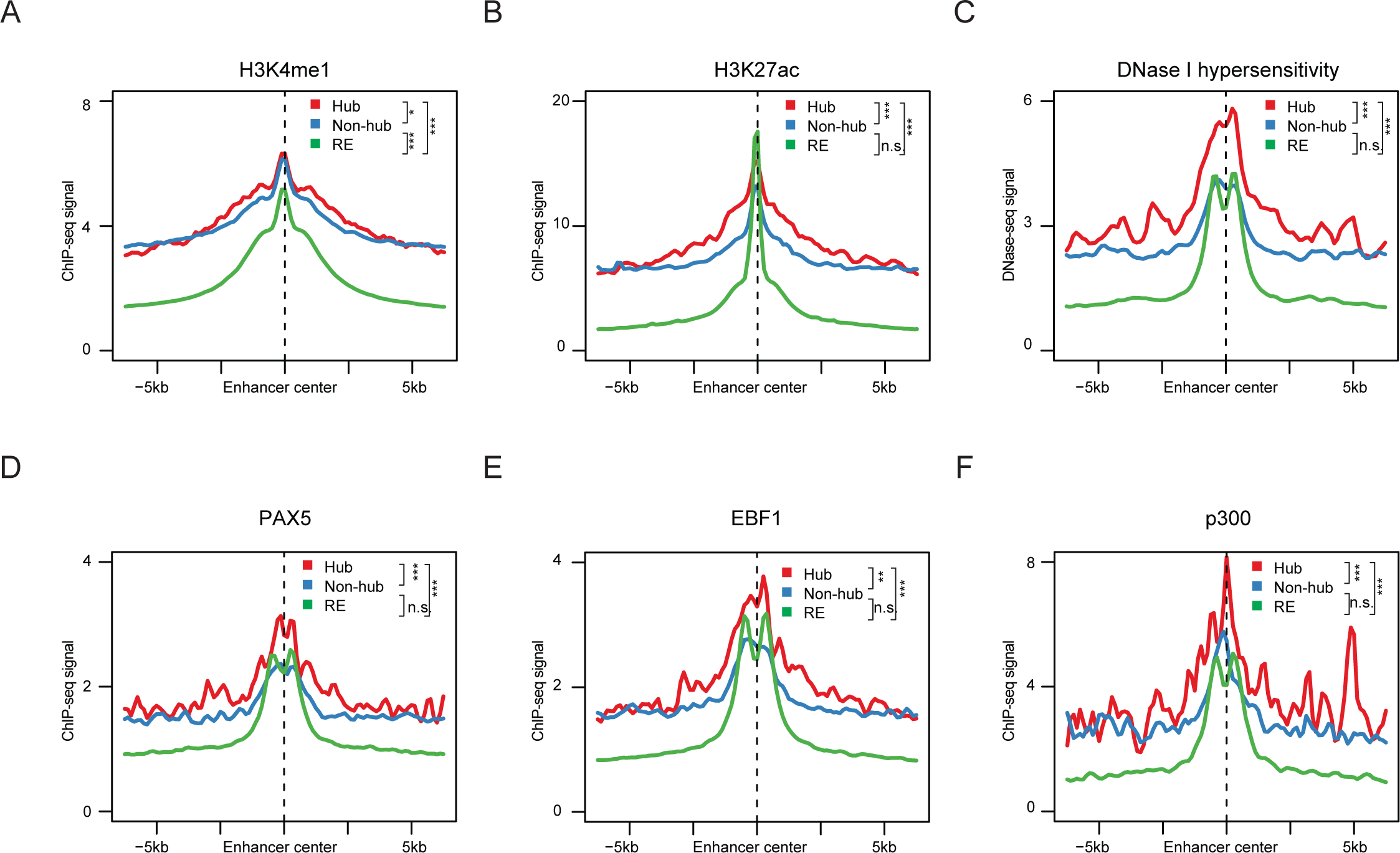
Chromatin landscapes around hub enhancers in GM12878 cells. (**A-F**) Spatial distribution of chromatin marks centered by enhancers in three groups, H3K4me1 (**A**), H3K27ac (**B**), DNase I hypersensitivity (**C**), master regulators PAX5 (**D**) and EBF1 (**E**), and coactivator p300 (**F**).

**Figure 3-figure supplement 1.**
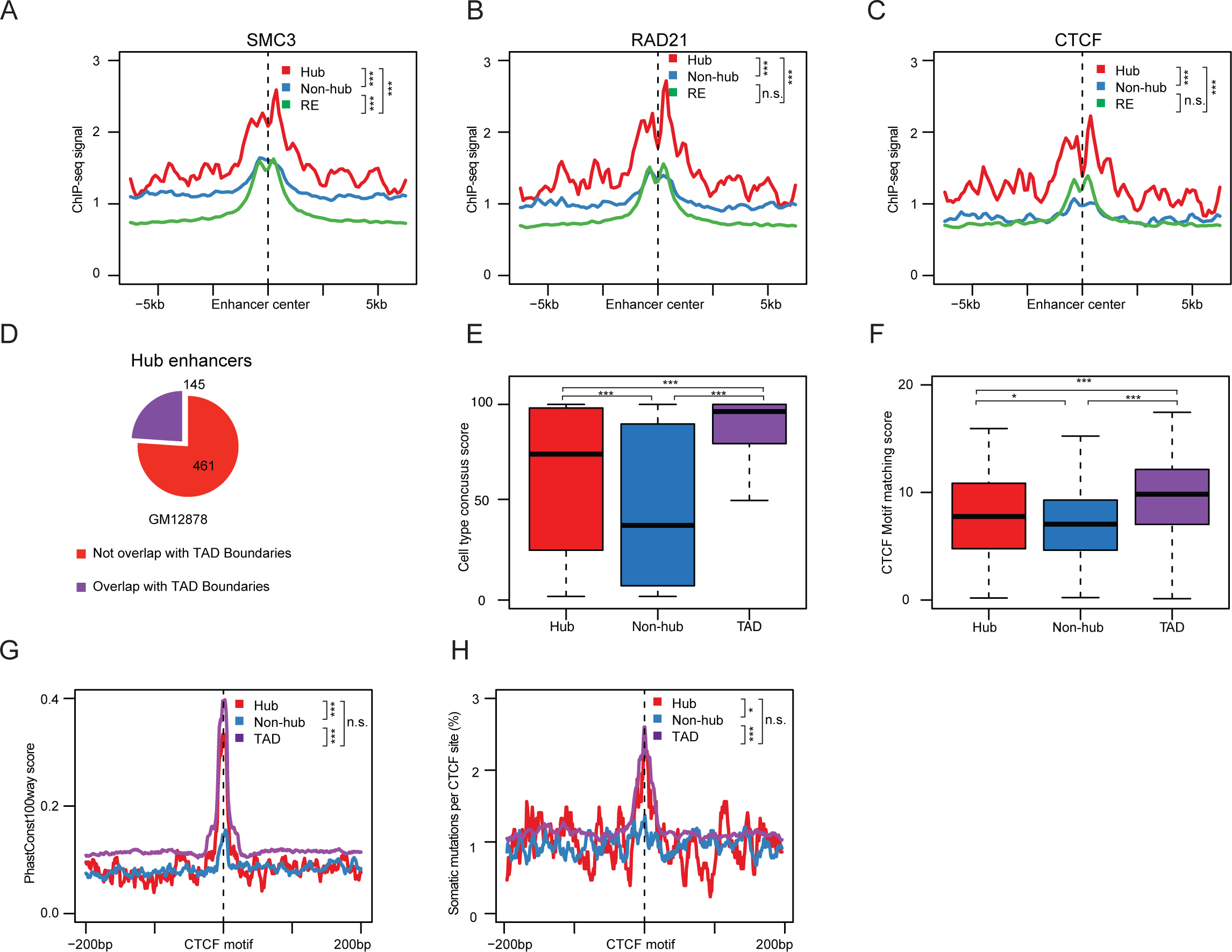
CTCF binding at hub enhancers within SEs hierarchy in GM12878 cells. (**A**-**C**) Spatial distribution of two cohesin components SMC3, RAD21 (**A**,**B**) and CTCF (**C**), centered by enhancers in three groups. (**D**) Percentage of hub enhancers with (purple) or without (red) overlapping with TAD boundaries collected from(Rao et al., 2014). The CTCF ChIP-seq peaks/motif-sites associated with hub enhancers overlapping with TAD boundaries were excluded for analysis in (**E-H**). (**E**) CTCF binding consensus across cell types in different contexts: hub (red), non-hub enhancers (blue) and TAD boundaries (purple). For each CTCF peak in GM12878, the consensus score (y-axis) was quantified as the percentage of cell types containing the same CTCF peak. *P* values are calculated using Student’s t test. \**P* < 0.05; \*\**P* < 0.01; \*\*\**P* < 0.001, n.s. not significant. (**F**) CTCF-motif-matching score (y-axis) of CTCF peaks. *P* values are calculated using Student’s t-test. \**P* < 0.05; \*\**P* < 0.01; \*\*\**P* < 0.001, n.s. not significant. (**G**) Sequence conservation around CTCF motif sites. The sitepro plots were centered by CTCF motif sites. *P* values are calculated using Student’s t-test based on the PhastConst100way score (y-axis) within CTCF motif sites. \**P* < 0.05; \*\**P* < 0.01; \*\*\**P* < 0.001, n.s. not significant. (**H**) Somatic mutation rate in cancers collected from IGGC around CTCF motif sites. The sitepro plots were centered by CTCF motif sites with 10bp smoothing window. *P* values are calculated using Fisher’s exact test based on overlap between CTCF motif sites and somatic mutation sites. \**P* < 0.05; \*\**P* < 0.01; \*\*\**P* < 0.001, n.s. not significant.

**Figure 4-figure supplement 1.**
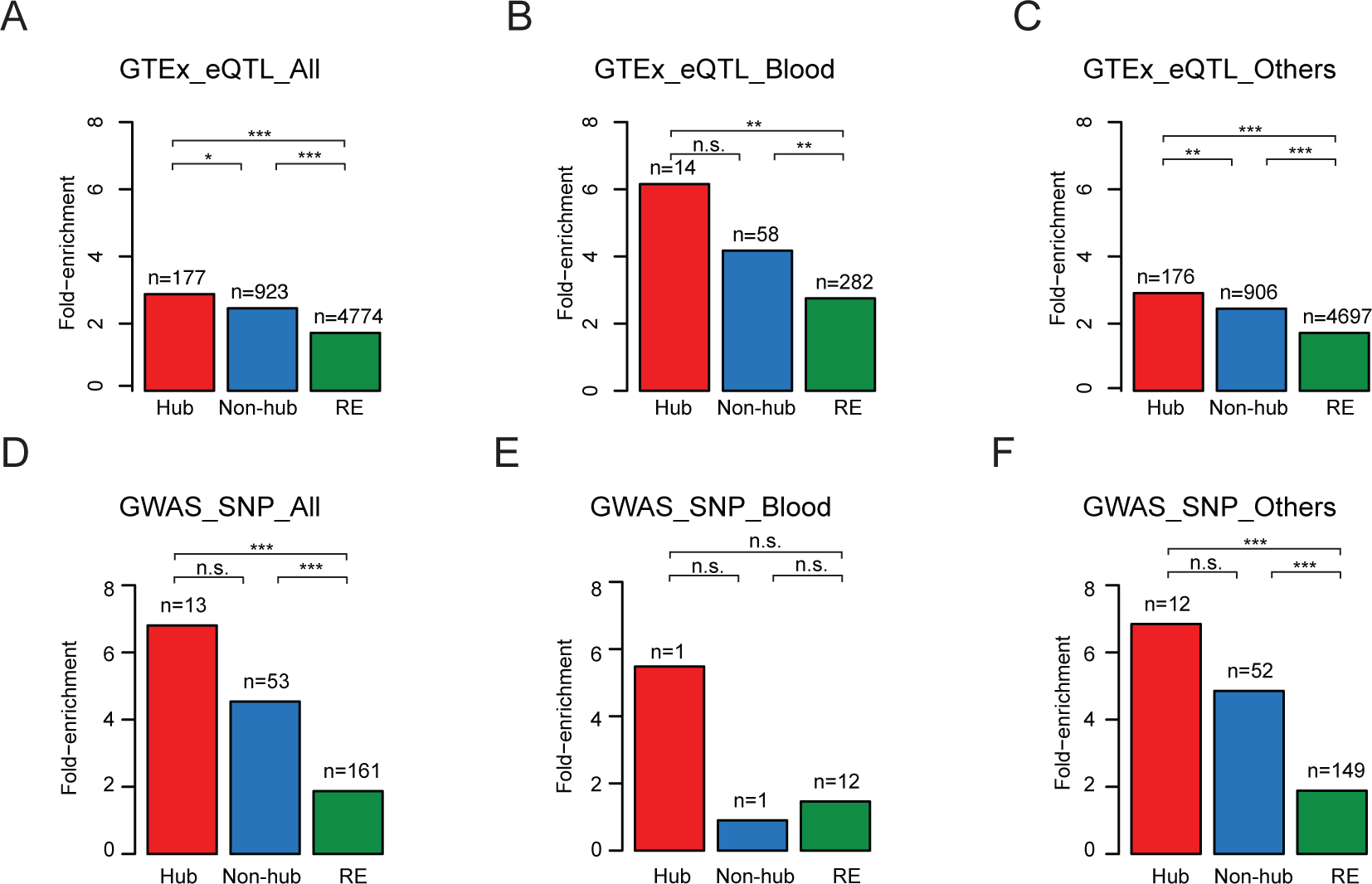
Enrichment of genetic variants associated with cell-type specific gene expression and diseases in hub enhancers in GM12878 cells. (**A-C**) Enrichment of the eQTLs curated in GTEx in the enhancers in three groups, using randomly selected genome regions as control (see Methods). The GTEx eQTL identified in all tissues (**A**) were separated into two subsets, identified in blood (**B**) or other tissues (**C**). The number of enhancers overlap in each group with eQTLs were labelled on each bar. *P* values are calculated using Fisher’s exact test. \**P* < 0.05; \*\**P* < 0.01; \*\*\**P* < 0.001, n.s. not significant. (**D-F**) Enrichment of the disease or traits-associated SNPs curated in GWAS catalog in the enhancers in three groups, using randomly selected genome regions as control (see Methods). The GWAS SNPs associated all diseases/traits (**D**), were separated into two subsets, associated with blood-related diseases/traits (**E**) or other traits (**F**). The number of enhancers overlap in each group with SNPs were labelled on each bar. *P* values are calculated using Fisher’s exact test. \**P* < 0.05; \*\**P* < 0.01; \*\*\**P* < 0.001, n.s. not significant.

**Figure 4-figure supplement 2.**
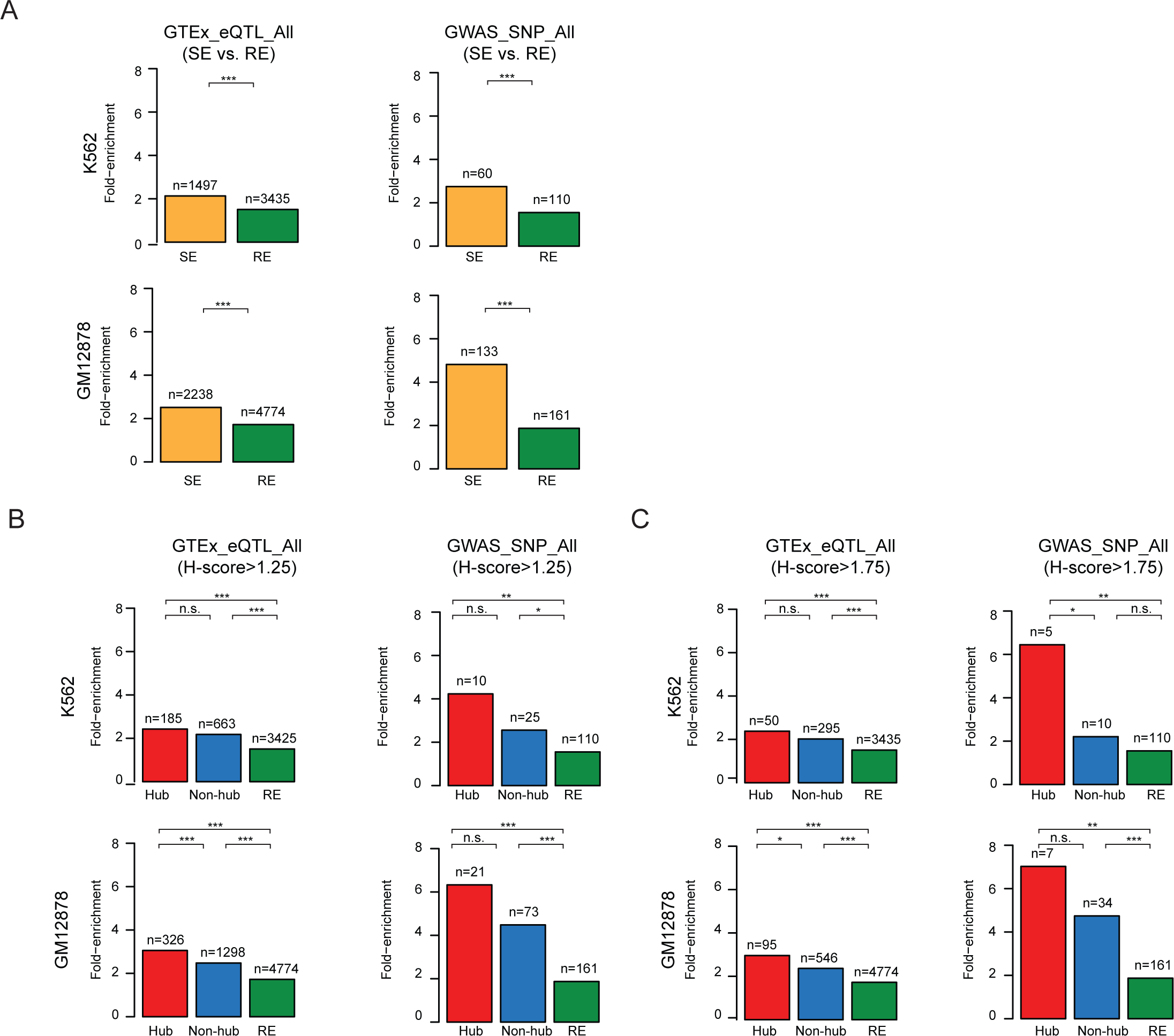
Enrichment of genetic variants associated with cell-type specific expression and diseases in K562 and GM12878. (**A**) Enrichment of GTEx eQTL (left) and GWAS SNPs (right) in SEs and REs in K562 (upper) and GM12878(lower). The number of enhancers overlap in each group with eQTLs were labelled on each bar. *P* values are calculated using Fisher’s exact test. \**P* < 0.05; \*\**P* < 0.01; \*\*\**P* < 0.001, n.s. not significant. (**B,C**) Enrichment of GTEx eQTL (left) and GWAS SNPs (right) in hub enhancers defined based on the threshold of H-score >1.25 (**B**) or H-score >1.75 (**C**) in K562 (upper) and GM12878(lower). The number of enhancers overlap in each group with eQTLs were labelled on each bar. *P* values are calculated using Fisher’s exact test. \**P* < 0.05; \*\**P* < 0.01; \*\*\**P* < 0.001, n.s. not significant.

**Figure 4-figure supplement 3.**
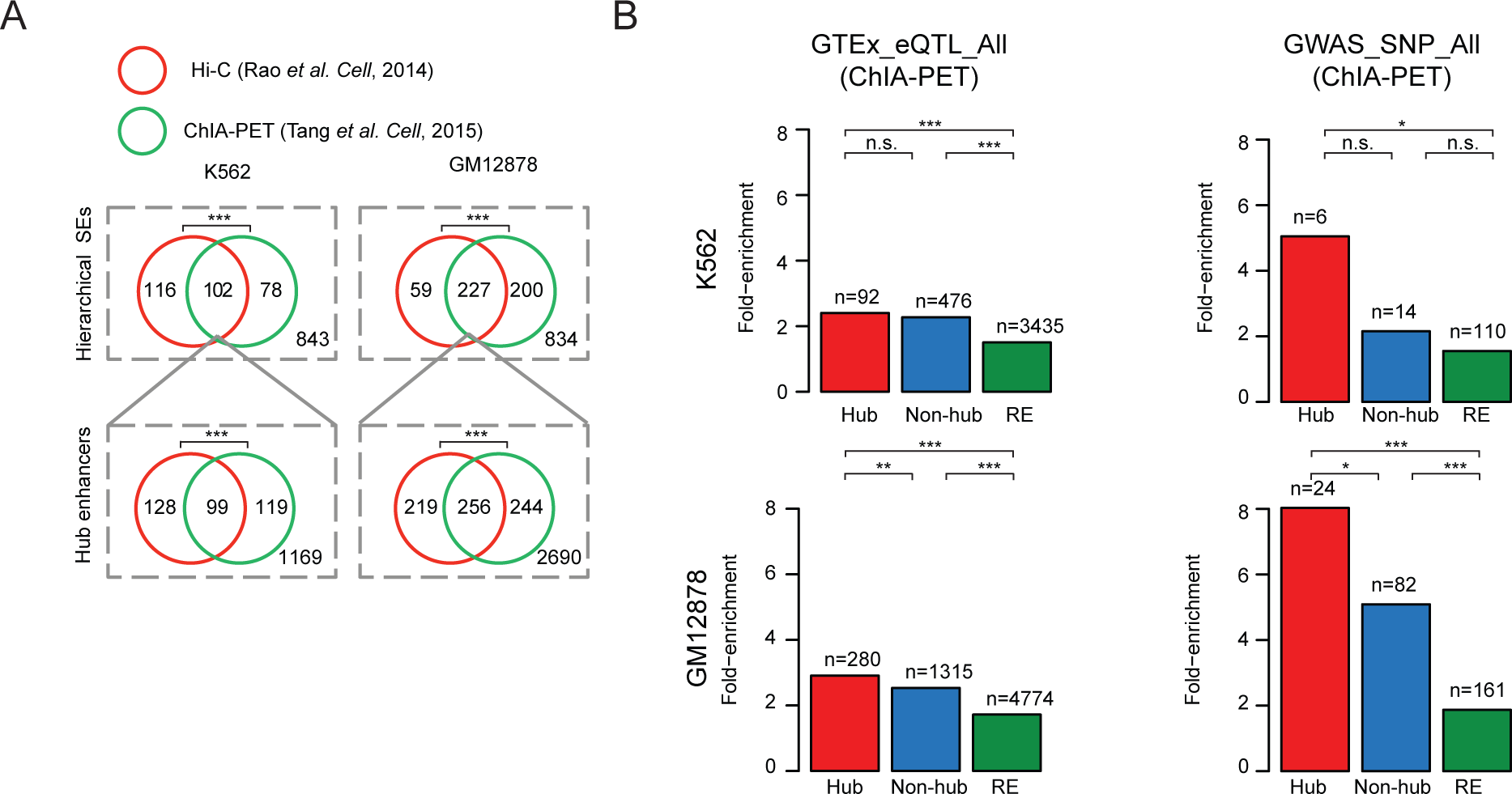
Comparison of hub enhancers defined based on chromatin interactions from Hi-C and ChIA-PET datasets in K562 and GM12878 cells. (**A**) Overlap between hierarchical SEs (left) or hub enhancers (right) using Hi-C and ChIA-PET dataset in K562 (upper) and GM1 2878 (lower). *P* values are calculated using Fisher’s exact test. \**P* < 0.05; \*\**P* < 0.01; \*\*\**P* < 0.001, n.s. not significant. (**B**) Enrichment of GTEx eQTL (left) or GWAS SNPs (right) in hub enhancers defined based on ChIA-PET. *P* values are calculated using Fisher’s exact test. \**P* < 0.05; \*\**P* < 0.01; \*\*\**P* < 0.001, n.s. not significant.

**Figure 5-figure supplement 1.**
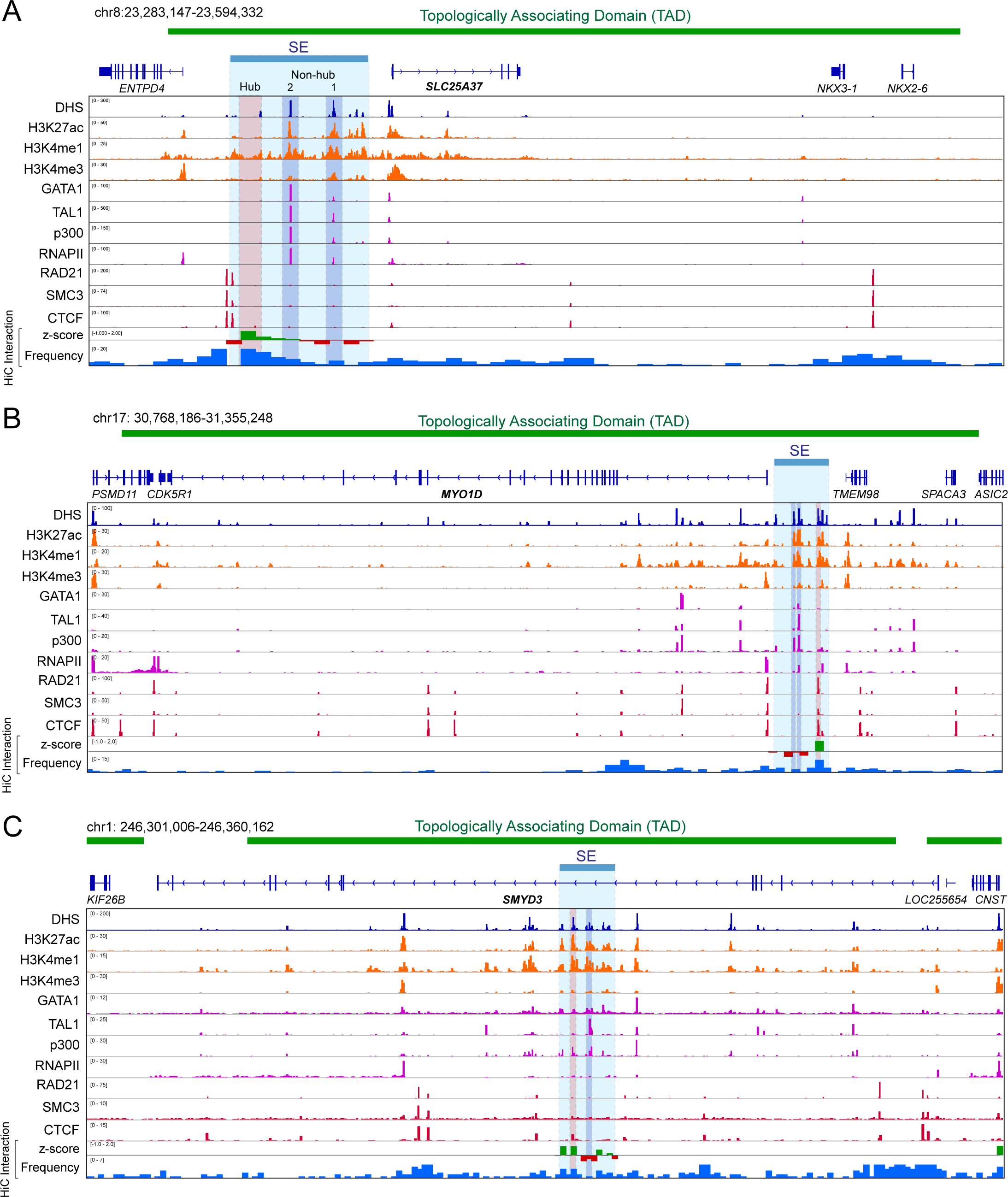
*In situ* analysis of the functional requirement of hub vs non-hub enhancers. (**A**) A genome browser view of the chromatin signatures and TF occupancy at the SLC25A37 SE locus in K562 cells. The identified SE is depicted by the blue shaded area. The hub and non-hub enhancers are denoted by the red and blue shaded lines, respectively. The Hi-C chromatin interaction z-score and frequency at 5kb resolution is shown at the bottom (see Methods). (**B**) A zoom-out view of the chromatin signatures and TF occupancy at the MYO1D SE locus in K562 cells is shown. (**C**) A zoom-out view of the chromatin signatures and TF occupancy at the SMYD3 SE locus in K562 cells is shown.

**Figure 5-figure supplement 2.**
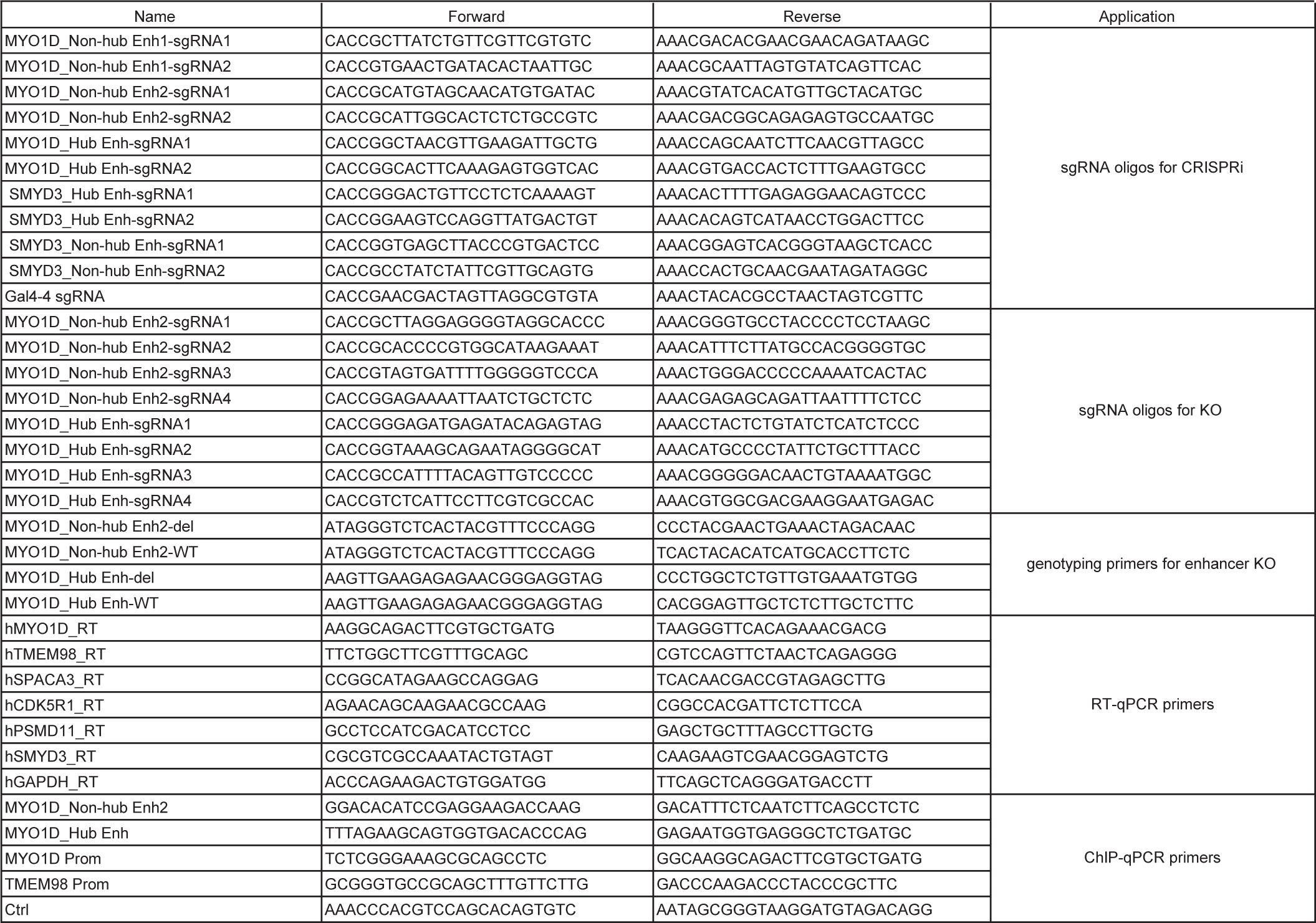
List of primer and sgRNA sequences used in this study.

